# Increased soluble amyloid-beta causes early aberrant brain network hypersynchronisation in a mature-onset mouse model of amyloidosis

**DOI:** 10.1101/723981

**Authors:** Inès R.H. Ben-Nejma, Aneta J. Keliris, Jasmijn Daans, Peter Ponsaerts, Marleen Verhoye, Annemie Van der Linden, Georgios A. Keliris

**Author notes:** ***Corresponding authors***, Inès R.H. Ben-Nejma, Bio-Imaging Lab, University of Antwerp, Universiteitsplein 1, 2610 Wilrijk, Antwerp Belgium, Prof. Dr. Georgios A. Keliris, Bio-Imaging Lab, University of Antwerp, Universiteitsplein 1, 2610 Wilrijk, Antwerp Belgium.

## Abstract

**Background:** Alzheimer’s disease (AD) is the most common form of dementia in the elderly population. Currently, no effective cure is available for AD. According to the amyloid hypothesis, the accumulation and deposition of the amyloid-beta (Aβ) peptides plays a key role in AD pathology. Soluble Aβ (sAβ) oligomers were shown to be synaptotoxic and involved in pathological hypersynchronisation of brain resting-state networks in different transgenic developmental-onset mouse models of amyloidosis. However, the impact of protein overexpression during brain postnatal development may cause additional phenotypes unrelated to AD. To address this concern, we investigated sAβ effects on functional resting-state networks in transgenic mature-onset amyloidosis Tet-Off APP (TG) mice.

**Methods:** TG mice and control littermates were raised on doxycycline (DOX) diet from 3d up to 3m of age to suppress transgenic Aβ production. Thereafter, longitudinal resting-state functional MRI was performed on a 9.4T MR-system starting from week 0 (3m old mice) up to 28w post DOX treatment. Ex vivo immunohistochemistry and ELISA analysis (additional mice cohort) was performed to address the development of amyloid pathology.

**Results:** Functional Connectivity (FC) analysis demonstrated early abnormal hypersynchronisation in the TG mice compared to the controls at 8w post DOX treatment. This effect was observed particularly across regions of the default mode-like network, known to be affected in AD. Ex vivo analyses performed at this time point confirmed a 20-fold increase in total sAβ levels and the absence of Aβ plaques in the TG mice compared to the controls. On the contrary at week 28, TG mice showed an overall hypoconnectivity, coinciding with a widespread deposition of Aβ plaques in the brain.

**Conclusions:** By preventing developmental influence of APP and/or sAβ during brain postnatal development, we demonstrated FC abnormalities driven by sAβ synaptotoxicity on resting state neuronal networks in mature-induced TG mice. Thus, the Tet-Off APP mouse model could be a powerful tool while used as a mature-onset model to shed light into amyloidosis mechanisms in AD. Therefore, this inducible APP expression model used in combination with early non-invasive in vivo rsfMRI readout for sAβ synaptotoxicity sets the stage for future Aβ targeting preventative treatment studies.

## BACKGROUND

Alzheimer’s disease (AD) is a devastating progressive neurodegenerative disorder, mainly characterized by the accumulation of amyloid-beta (Aβ) plaques and neurofibrillary tangles (NFTs), which leads to dementia [1]. Most of AD patients develop the late-onset sporadic form of AD (sAD), while the early-onset familial form of AD (fAD) is rare (<1% of AD cases) [2–4]. Ageing has been identified as the greatest known risk factor of sAD, with the prevalence doubling every 5 years for people over 65 and reaching nearly one third by the age of 85 [1]. As the ageing population is increasing, the number of AD patients suffering from dementia will raise considerably [5].

Evidence from studies on fAD (caused by genetic mutations) supported the idea, referred as “the amyloid hypothesis”, that the abnormal accumulation of extracellular soluble Aβ (sAβ) peptides is initiating AD pathology [6, 7]. Under normal conditions, sAβ peptides are naturally generated from the proteolysis of the amyloid precursor protein (APP), which is a single-pass transmembrane protein expressed at high levels in the brain and concentrated in the synapses of neurons [8]. Further, sAβ monomers are thought to have a physiological role in controlling synaptic activity, excitability and cell survival [9]. Under pathological conditions and upon ageing, a metabolic dysregulation seems to cause the accumulation of sAβ peptides in the extracellular space, which oligomerize and aggregate, forming insoluble Aβ plaques [10], [11]. It is believed that sAβ oligomers are more synaptotoxic than the plaques, altering synaptic transmission and causing synapse loss and neuronal death [2, 12, 13], mainly in the cerebral cortex and certain subcortical regions [14], resulting in a progressive loss of cognitive functions [11].

Despite the important efforts to develop therapeutics, there is currently no cure available for AD. As the first symptoms of AD appear decades after the onset of the disease, scientists have moved their focus on the early stage of the disease, including both therapeutic treatments and early diagnosis. With the evolution of neuroimaging techniques, some functional AD-related alterations have been highlighted, particularly using resting-state functional magnetic resonance imaging (rsfMRI). This in vivo non-invasive imaging technique measures the blood oxygen level-dependent (BOLD) low-frequency signal fluctuations of neuronal activity to indirectly evaluate brain network functions [15, 16]. The temporal correlation of these fluctuations between spatially distinct brain regions has been suggested to reflect the brain’s functional connectivity (FC). Therefore, using rsfMRI, we can detect alterations at the neuronal network level, reflecting impaired communication between brain regions caused by synaptic dysfunction. In addition, it has been demonstrated that AD patients and individuals with mild cognitive impairment displayed disturbed FC in specific brain networks, such as the default mode network (DMN) [17–20]. The DMN was identified as a network of anatomically distant brain regions which show highly correlated synaptic activity when a person is at rest [21, 22]. Thus, rsfMRI performed at early stages can provide important information on the FC and its potential disturbances, crucial for early diagnosis of AD.

Increasing evidence towards oligomeric sAβ as an early biomarker, instead of plaques that appear later in the course of disease pathology, made the research focus on the role of sAβ oligomers in AD development. For instance, synaptic transmission and plasticity were altered months before plaque deposition in the hippocampus of young AD transgenic mice [23]. Behavioural deficits were also found in the pre-plaque stage [24]. In 2012, Busche et al. showed, using *in vivo* two-photon Ca2+ imaging, that sAβ oligomers caused very early functional impairment in AD transgenic mice due to hippocampal hyperactivity [25]. Moreover, they also reported that sAβ oligomers altered the slow-wave propagation, causing a long-range slow-wave activity breakdown [26]. Interestingly, rsfMRI performed in different transgenic mouse models of AD initially reported decreased FC at a stage of abundant plaque deposition [27, 28]. As the research focus moved from insoluble amyloid deposits towards oligomeric sAβ, rsfMRI performed before plaque deposition revealed an early pathological hypersynchronisation of brain resting state networks, especially in the DMN [29]. This evidence of oligomeric sAβ-induced synaptotoxicity has, however, come directly from transgenic developmental-onset mouse models of amyloidosis, i.e. the overexpression of APP starts from embryonic or very early postnatal developmental stage. As overexpression, and consequently, overproduction of APP and Aβ, occurs during brain development, this might affect the cell signalling and synapse formation, causing artificial phenotypes unrelated to AD pathology [4, 30]. Moreover, immature and mature mice might respond differently to the high exposure of sAβ species, making the neuronal circuits more sensitive or resilient to sAβ [31]. Therefore, it is crucial to evaluate the toxicity of sAβ oligomers on mature mice. However, all available transgenic models of amyloidosis contain the fAD mutations, thus modelling the amyloid pathology of fAD (developmental-onset) and not sAD [4].

In order to evaluate the impact of oligomeric sAβ accumulation on the brain neuronal networks in mature mice, we used the inducible Tet-Off APP mouse model of amyloidosis (line 107) in which the APP overexpression can be controlled via a specific diet [32]. APP overexpression was switched-on when the mice were 3m of age, when the brain can be considered as mature [33]. As the most drastic developmental changes in both myelination and cortical thickness have already taken by that age, this model was referred as mature-onset APP expression, which is in a context resembling closer the human sAD amyloid pathology. Using rsfMRI and ex vivo brain analyses, we investigated the toxicity of increased sAβ oligomers in mature-onset Tet-Off APP mice on brain neuronal networks.

## METHODS

### Animals

The Tet-Off APP transgenic mouse model of amyloidosis (tetO-APPswe/ind line 107 [32]) used in this study allows the controllable expression of a chimeric mouse APP containing a humanized Aβ domain (mo/huAPP695) using the Tet-Off system. Single transgenic tTA and APP males were kindly provided by Prof. Dr JoAnne McLaurin (Sunnybrook Health Sciences Centre, Toronto, Canada). The single transgenic males were crossed with non-transgenic females on a C57BL6/J background (Charles River, France) to establish and maintain the single transgenic colonies. The bigenic (TG) animals were obtained by crossing APP mice, in which a tetracycline-responsive (tetO) promoter drives the expression of the APP transgene bearing the Swedish and Indiana mutations (APPswe/ind), with transgenic mice expressing the tetracycline-Transactivator (tTA) gene. As tTA is under control of the CaMKIIα promoter, APP expression is neuron-specific, with moderate levels (about 10 fold higher than endogenous mouse APP) and mainly localized in the forebrain [32]. Once the APP expression is turned on, the amyloid plaques start to deposit approximately after 2m [32, 34].

### Doxycycline treatment

The APP expression is reversibly turned on or off in the absence or presence of the tetracycline (antibiotic) or derivatives (here, doxycycline - DOX), respectively (Tet-Off expression system, Additional Figure 1). APP expression was therefore inhibited by feeding females with litters and weaned pups with DOX diet (100 mg/kg Doxycycline diet; Envigo RMS B.V., The Netherlands). DOX treatment started from 3d postnatal to avoid epigenetic reduction in transgene expression [35] up to 3m of age and switched to a regular chow (referred as week 0) until the end of the longitudinal experiment (referred as week 28). As the cortex keeps on flattening during the first 3m and most drastic changes in myelination have already taken place by that age, the brain is considered as ‘mature’ by 3m of age [33].

**Additional Figure 1.**
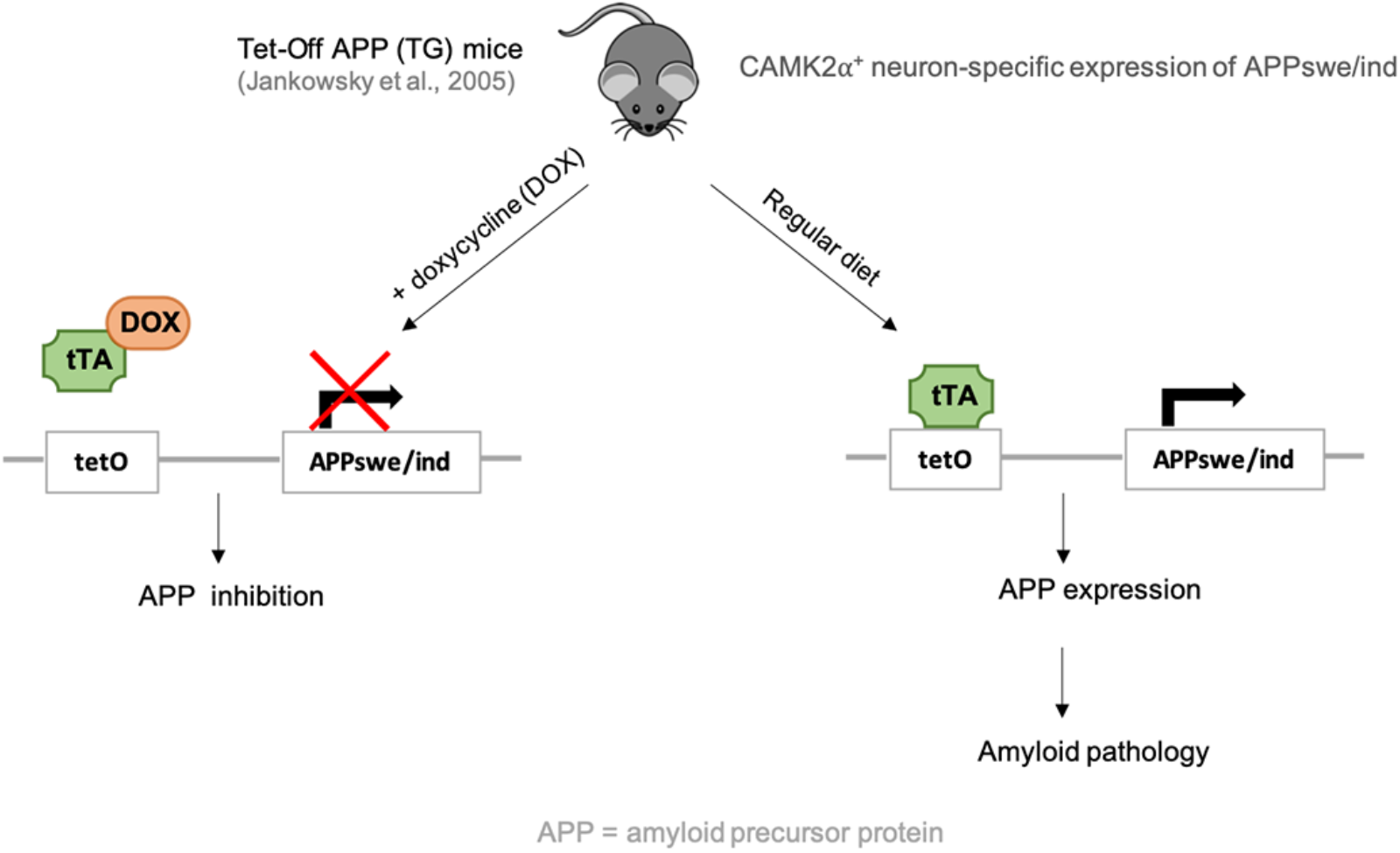
Tet-Off APP mouse model. The Tet-Off APP mice, bearing a chimeric APP (mo/huAPP695) which is co-expressed with the tetracycline-Transactivator (tTA). The APP expression can be turned off (left panel) or on (right panel) in the presence or absence of doxycycline (DOX) respectively.

### Genotyping

Transgenic animals and control littermates were genotyped from ear cuts. First, a DNA extraction was performed (QIAamp DNA Mini Kit (250), QIAGEN Benelux B.V., Belgium), followed by two separate PCR reaction with Extract-N-Amp™ PCR ReadyMix (Sigma-Aldricht, Belgium) and specific primers (all from Invitrogen, ThermoFisher Scientific, Belgium) as described previously [32]. The PCR reaction mixture was incubated at 94°C for 3 min, then 40 cycles of amplification (94°C, 30 s; 60°C (tTA) or 55°C (APP), 1 min; 72°C, 1 min), and finally elongation at 72°C for 4 min.

### Animal handling

Three groups were subjected to a longitudinal MRI study: a TG group, a tTA group (control for promoter-induced changes) and a wild-type (WT) group (control for DOX-induced changes). Mice (N=37, mixed in gender) were scanned at week 0, 8, 16 and 28 after termination of DOX diet. Five animals (4 TG and 1 WT) were excluded from the study due to unexpected death (3 TG before week 8, and 1 TG and 1 WT after week 8 post DOX treatment). All MRI experiments were performed on spontaneously breathing mice under isoflurane anaesthesia (3% for induction, 2% for animal positioning in the bed and maintenance, Isoflo®, Abbot Laboratories Ltd., Illinois, USA), administered in a gaseous mixture of 33% oxygen (200 cc/min) and 66% nitrogen (400 cc/min). To have a reproducible flat-skull position during the MRI experiments, the head of the animals was immobilized in an MRI compatible mouse stereotactic device, including a nose cone in which the anaesthetic gas was delivered, with a tooth bar and blunt earplugs. For the rsfMRI acquisition, a bolus of 0.05 mg/kg with medetomidine (Domitor, Pfizer, Karlsruhe, Germany) was administered subcutaneously (s.c.) and isoflurane was decreased immediately to 1%. Ten minutes after the bolus injection, a continuous infusion of medetomidine (0.1 mg/kg/h) was started and 10 min before the rsfMRI acquisition, isoflurane was decreased and kept to 0.4% to maintain an optimal sedation level for the 10 min acquisition of the rsfMRI data [36, 37]. Afterwards, isoflurane levels were gradually increased to 2% within the 10 next min, according to the respiration of the animals, to keep them stable during the acquisition of anatomical 3D scans. To counteract the effect of medetomidine, a subcutaneous injection of 0.1 mg/kg atipamezole (Antisedan, Pfizer, Karlsruhe, Germany) was administered after the MRI procedures. During the imaging experiments, the physiological status of all animals was closely monitored. The respiration rate was monitored with a small animal pressure sensitive pad (MR-compatible Small Animal Monitoring and Gating System, SA instruments, Inc.) and maintained within normal physiological range. Rectal temperature was maintained at (37.0 ± 0.5) °C using a feedback controlled warm air system (MR-compatible Small Animal Heating System, SA Instruments, Inc.). A fibre optic pulse oximetry sensor (SA instruments, Inc.) positioned on the tail of the mice was used to monitor blood oxygenation.

### MRI acquisition

The imaging measurements were performed on a 9.4 T Biospec MRI system (Bruker BioSpin, Germany) with the Paravision 6.0 software (www.bruker.com) using a standard Bruker cross coil set-up with a quadrature volume coil and a 2×2 surface array mouse head receiver coil. Three orthogonal multi-slice Turbo RARE T2-weighted images were acquired to ensure uniform slice positioning (RARE; TR/TE 2500/33 ms; 9 slices of 0.7 mm; FOV (20×20) mm^2^; pixel dimensions (0.078×0.078) mm^2^. A field map was acquired for each animal to assess field homogeneity, followed by local shimming, which corrects for the measured inhomogeneity in an ellipsoid volume of interest (VOI) within the brain. RsfMRI acquisition was performed using a multi-slice T2*-weighted single shot gradient echo (GE) echo planar imaging (EPI) sequence (2D GE-EPI; TR/TE 2000/17 ms; 12 horizontal slices of 0.4 mm, slice gap 0.1 mm; 300 repetitions). The field-of-view (FOV) was (20×20) mm^2^ and the matrix size (96×64), resulting in voxel dimensions of (0.208×0.313×0.5) mm^3^. The acquisition of the rsfMRI scans started 40 min after the bolus injection and the total scan time was 10 min. Anatomical 3D images were acquired using a 3D RARE sequence (TR/TE 1800/42 ms; RARE factor 16). The reconstructed matrix size was (256×256×128) and FOV (20×20×10) mm^3^, resulting in a voxel resolution of (0.078×0.078×0.078) mm^3^ with a total scan duration of 44 min.

### MRI data pre-processing

Pre-processing of the rsfMRI data was performed using SPM12 software (Statistical Parametric Mapping, http://www.fil.ion.ucl.ac.uk) in MATLAB (MATLAB R2016a, The MathWorks Inc. Natick, MA, USA) and included realignment, co-registration, normalization and spatial smoothing, as described previously [29, 37]. Briefly, all rsfMRI images within each session were first realigned to the first image, using a least-squares approach and a 6-parameter rigid body spatial transformation. Then, rsfMRI images were co-registered to the subject’s 3D RARE scan acquired during the same imaging session using mutual information as the similarity metric. In parallel, an in-house 3D-template, based from 3D RARE images of similar mice aged from 3m to 5m old was created with Advanced Normalization Tools (ANTs), using a global 12-parameter affine transformation followed by a nonlinear deformation protocol. This template was used to spatially normalize each 3D RARE scan. Afterwards, all of the transformations (realignment, co-registration and normalisation) were applied during the spatial normalization step. Lastly, the rsfMRI data was smoothed in-plane, using a Gaussian kernel with full width at half maximum of twice the voxel size (FWHM = 0.416 mm x 0.625 mm). Finally, the rsfMRI data was filtered between 0.01 and 0.1 Hz using the Resting State fMRI Data Analysis toolbox (REST 1.8, http://resting-fmri.sourceforge.net/).

### MRI data analysis

First, a group Independent Component Analysis (ICA) was performed to determine which brain resting-state networks could be detected, as described previously [29, 37]. This analysis was done within the population, including all data sets (3 groups, 4 time points), using the GIFT-toolbox (Group ICA of fMRI toolbox, version 3.0a: http://icatb.sourceforge.net/), with a pre-set of 20 components. The identification of relevant neuronal components was based on a 3D anatomical template and with reference to the 3^rd^ edition of the anatomical mouse brain atlas from Paxinos and Franklin. For each component, corresponding to a specific brain region, a region-of-interest (ROI) within the component was manually delineated in each hemisphere separately, using MRIcron software (https://www.nitrc.org/projects/mricron/) with the ellipse shape tool of 31 voxels. These ROIs were used for correlation analyses, where pairwise correlation coefficients between each pair of ROIs were calculated and z-transformed using an in-house program developed in MATLAB (MATLAB R2014a, The MathWorks Inc. Natick, MA, USA). Mean z-transformed functional connectivity (FC) matrices were calculated for each group at each time point.

### Immunohistochemistry

An additional group of mice was used for histology. Brain samples were collected at 8w (N=2/group; actual age: 5m) and 28w post DOX treatment (N_WT_=1; N_TG_=1; actual age 10m) as described previously [29]. Briefly, mice were deeply anesthetized following an intraperitoneal injection of 60 mg/ kg/BW pentobarbital (Nembutal; Ceva Sante Animale, Brussels, Belgium), after which a transcardial perfusion first with ice-cold PBS and then with 4% paraformaldehyde (Merck Millipore, Merck KGaA, Darmstadt, Germany) was performed. Next, brains were surgically removed and post-fixed in 4% paraformaldehyde for 2h. To freeze protect the fixed brains, a sucrose gradient (sucrose, Sigma Aldrich) was applied: 2h at 5%, 2h at 10%, and overnight at 20%. Subsequently, brains were snap frozen in liquid nitrogen and stored at −80°C until further processing. Ten-μm sagittal sections were prepared using a cryostat (CryoStar NX70; ThermoScientific). For immunofluorescence analyses, the following primary antibodies were used: chicken anti GFAP (Abcam ab4674) 1:1000, rabbit anti IBA-1 (Wako 019-19741) 1:1000, and the following secondary antibodies: donkey anti chicken (Jackson 703-166-155) 1:1000, goat anti rabbit (Jackson 111-096-114) 1:1000. Furthermore, TOPRO staining (Life technologies T3605, 1:200) and thioflavin-S staining (Santa Cruz Biotechnology, sc-215969) were performed to, respectively, stain the nuclei and to visualize amyloid plaques. Following staining, sections were mounted using Prolong Gold Antifade (P36930; Invitrogen). All images were acquired using an Olympus BX51 fluorescence microscope equipped with an Olympus DP71 digital camera. Image acquisition was done with CellSens Software (Olympus, Tokyo, Japan, http://www.olympus-global.com). To localize the regions of interest, the mouse brain atlas from Paxinos and Franklin was used as reference. Images were further processed with ImageJ Software 1.52k (National Institutes of Health, https://imagej.nih.gov/ij/download.html).

### Enzyme-linked immunosorbent assay (ELISA)

An ELISA analysis of total sAβ_1-x_ was performed on hemibrains of an extra group of TG mice and control littermates (N=3/group) at 8w post DOX treatment, adapted from [29]. Briefly, the mice were deeply anesthetized with an intraperitoneal injection of 60 mg/ kg/BW pentobarbital (Nembutal; Ceva Sante Animale, Brussels, Belgium), followed by a transcardial perfusion with ice-cold PBS. Then, the brains samples were surgically removed and the hemispheres were split. Both hemispheres were snap frozen in liquid nitrogen and stored in separate tubes at −80°C. Next, to extract the sAβ fraction, samples (100mg tissue/mL) were homogenized in a 0.2% diethylamine solution (DEA, 0.2% in 50mM NaCl), using a motor-driven tissue grinder for 30 s. Then, the samples were centrifuged at 20 800 g for 90 min at 4°C to clarify the homogenate. The next step consisted in collecting the supernatant and centrifuging it at 20 800 g for 20 min at 4°C. As the sAβ fraction was restrained to the resulting supernatant, the latter was collected and immediately neutralized by adding 1/10 volume 0.5 M Tris HCl (pH 6.8). The samples for ELISA were stored at −80°C. Finally, sandwich ELISA (IBL International GMBH, 27729) was performed to determine the levels of sAβ_1-x_, according to the manufacturer’s guidelines. The ELISA signal was quantified using a microplate reader (EnVision 2103 Multilabel Reader, PerkinElmer).

### Statistical analyses

All statistical analyses were first performed including the three groups (WT, tTA and TG). As no significant differences were found between the two control groups (i.e. WT and tTA), these control genotypes were treated equally and the data from these two groups was collapsed as one control group (Ctrl) [31].

ROI-based analysis was assessed using a linear mixed model in JMP Pro 14.0, with Time, Genotype, the interaction [Time*Genotype] as fixed effect and subject as random effect. Tukey’s HSD (honestly significant difference) test with p<0.05 was used as *post hoc* test. A false discovery rate (FDR) correction (p<0.05) was applied to account for the multiple testing, while performing multiple analyses in parallel. GraphPad Prism was used to make the graphs. All results are shown as mean ± SEM (standard error of mean).

For *ex vivo* data, ELISA test was analysed in JMP Pro 14.0 using a non-parametric Wilcoxon test to test for the main effect of genotype at 8w post DOX treatment.

## RESULTS

### TG mice presented an alteration of FC over time

Independent component analysis (ICA) that was performed across the population revealed eleven anatomically relevant neuronal components in the cortex and subcortical regions which were further clustered into larger functional networks such as the DMN, sensory and subcortical networks (Additional Figure 2).

**Additional Figure 2.**
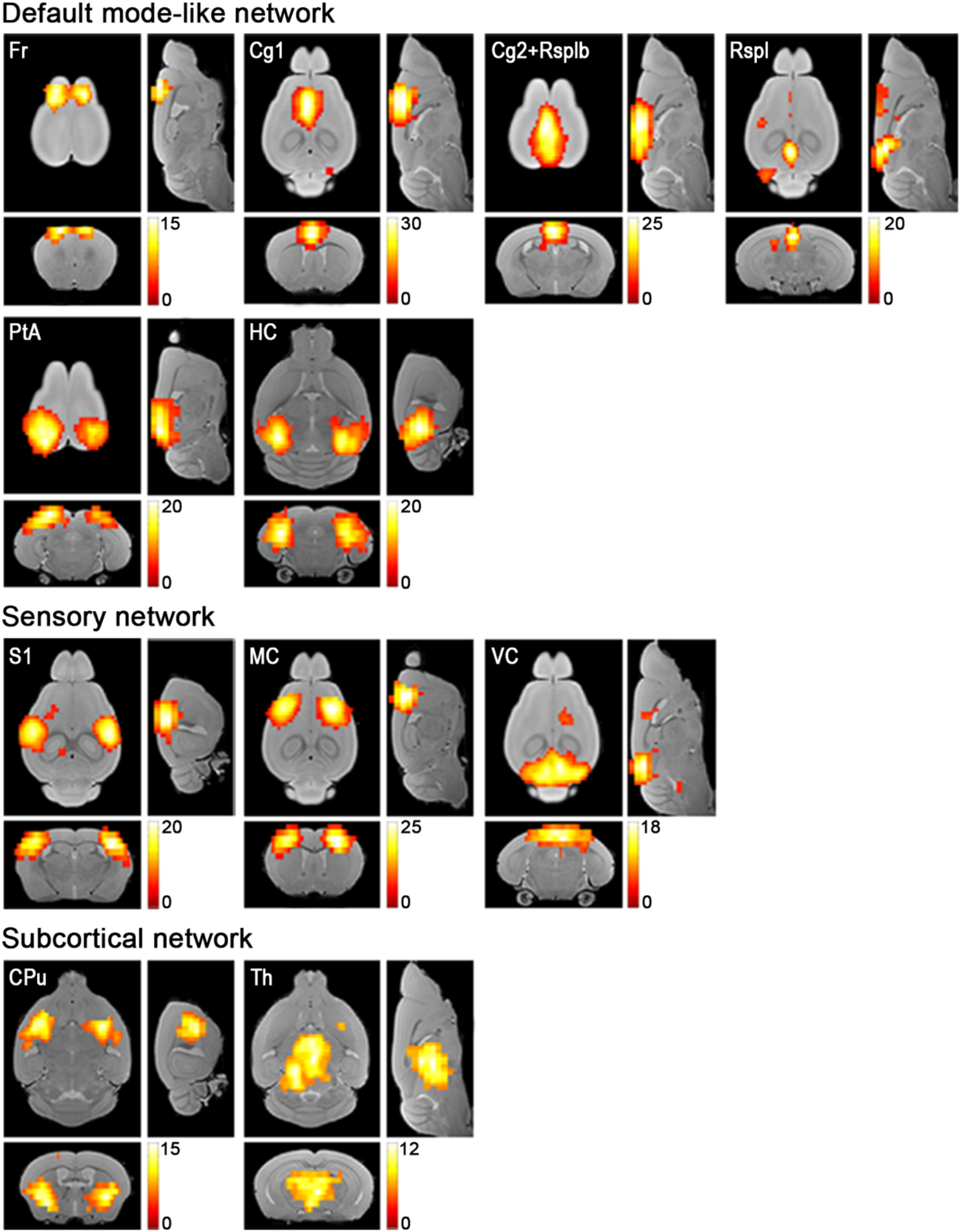
Overview of the ICA components of interest. Statistical maps are overlaid on an anatomical 3D template. The colour scale indicates t-values, with yellow representing stronger correlation. An FWE correction of p<0.05 and a minimum cluster size of 10 voxels was applied. Abbreviations: see **Table 1**.

The naming of each component was chosen based on the main anatomical brain regions included in the component (Table 1). The remaining components consisted of some duplicates (one bilateral and two unilateral in the left and right primary somatosensory cortex), three were localized in the cerebellum, one in the brainstem and two non-neuronal components in the ventricles; these components were considered of no interest and were not further analysed.

**Table 1.** List of the main regions-of-interest (ROI) defined guided by the ICA components.

| ICA components | Abbreviations |
| --- | --- |
| Hippocampus | LHC ; RHC |
| Cingulate (anterior part) | LCg1 ; RCg1 |
| Cingulate (posterior part) + Retrosplenial (anterior part) | LCg2+RSplb ; RCg2+RSplb |
| Retrosplenial (posterior part) | LRSpl ; RRSpl |
| Parietal association area | LPtA ; RPtA |
| Frontal area | LFr ; RFr |
| Somatosensory area | LS1 ; RS1 |
| Motor area | LMC ; RMC |
| Visual cortex | LVC ; RVC |
| Caudate putamen | LCPu ; RCPu |
| Thalamus | LTh ; RTh |

In order to assess the functional connectivity per group at each time point, we performed region-of-interest (ROI)-based analysis. To this end, ROIs were delineated based on stereotactic coordinates and guided by the eleven neuronal components of interest, which were used as a template. The resulting connectivity matrices combined for the TG (sub-diagonal) and Ctrl (supra-diagonal) groups are presented for week 0, 8, 16 and 28 post-DOX treatment (Figure 1A-D).

**Figure 1.**
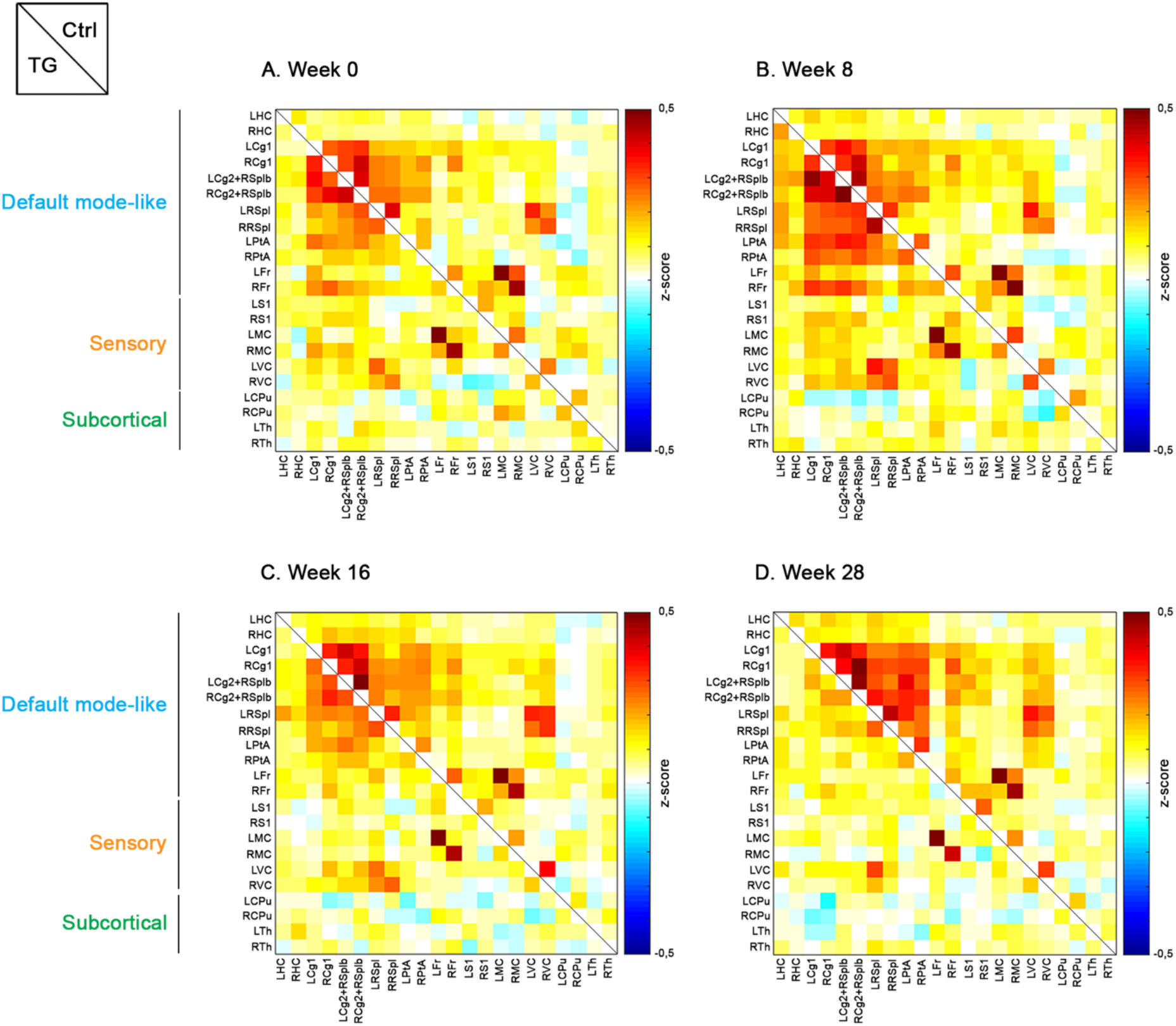
ROI-based functional connectivity analysis. Functional connectivity matrices show the average z-transformed functional connectivity (zFC) for Ctrl (supra-diagonal) and TG (sub-diagonal) animals at week 0 (**A**), 8 (**B**), 16 (**C**) and 28 (**D**) post DOX treatment. Each square indicates the zFC between a pair of ROIs. The colour scale represents the connectivity strength with white indicating a low zFC and red/blue indicating positive/negative zFC values.

Notably, changes including both increases and decreases in FC were observed across both groups and time. To investigate the dynamics of FC changes, we first examined the average FC across the whole brain (i.e. average of all ROIs). The FC demonstrated a trend (p=0.07) for increased FC in TG animals relative to controls 8w post DOX treatment (pre-plaque stage); FC was then gradually decreased over time resulting to a significantly reduced FC (p=0.0004) 28w post-DOX treatment (post-plaque stage, Figure 2A).

**Figure 2.**
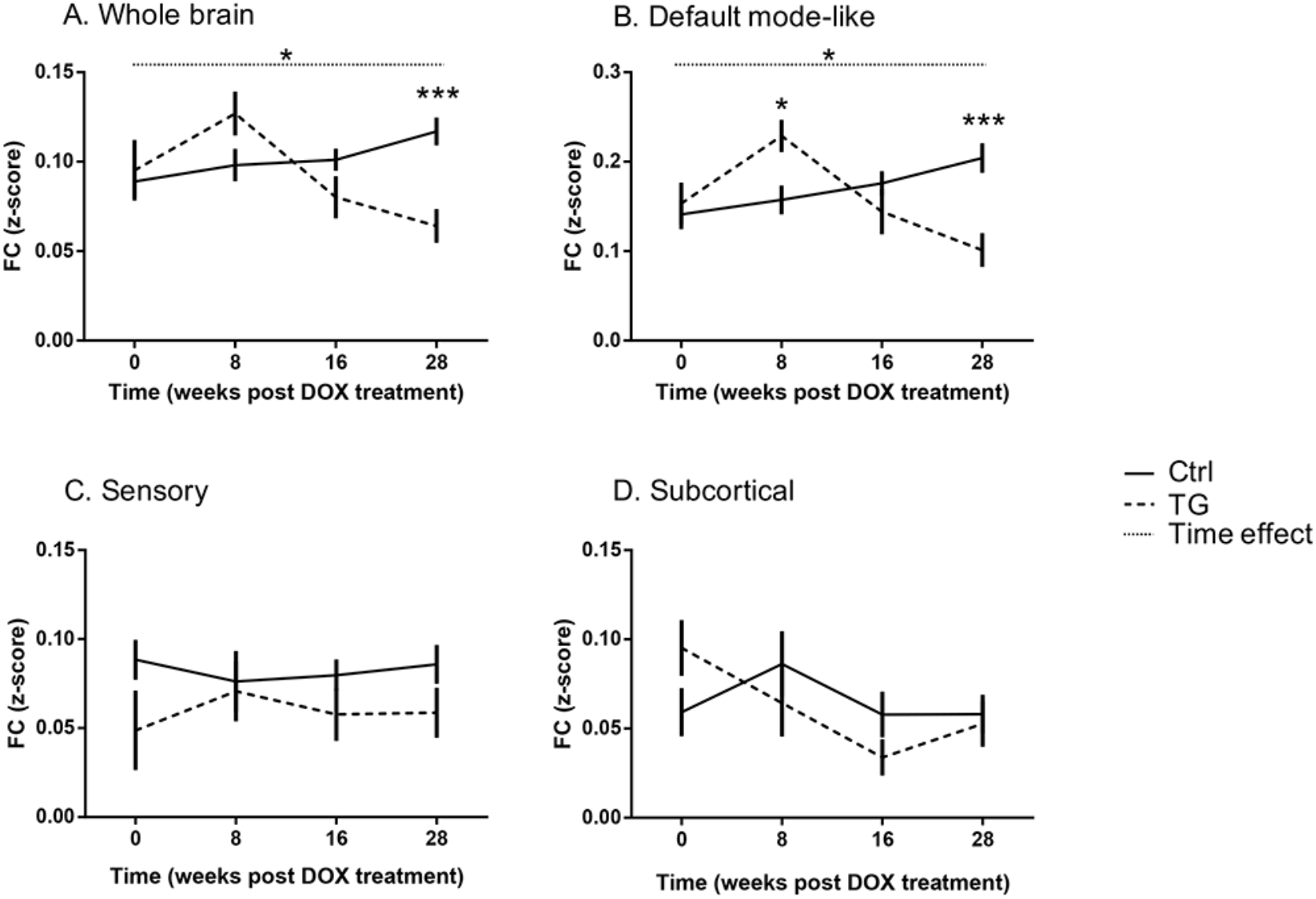
Average FC within each network. Mean FC (z-score) over time for both groups in the whole brain (**A**), the default-mode like network (**B**), the sensory network (**C**) and the subcortical regions (**D**). All values are presented as mean ± SEM; dashed line corresponds to TG group and full line to the Ctrl group. Significant [Genotype*Time] interaction and Time effect are indicated with stars and dotted line, respectively. *p<0.05; ***p<0.001.

The outcome of the statistical analyses is summarized in Table 2. To further understand which networks contributed to this pattern of connectivity, we then performed similar analysis restricted within three networks (DMN, Sensory, and subcortical).

**Table 2.**
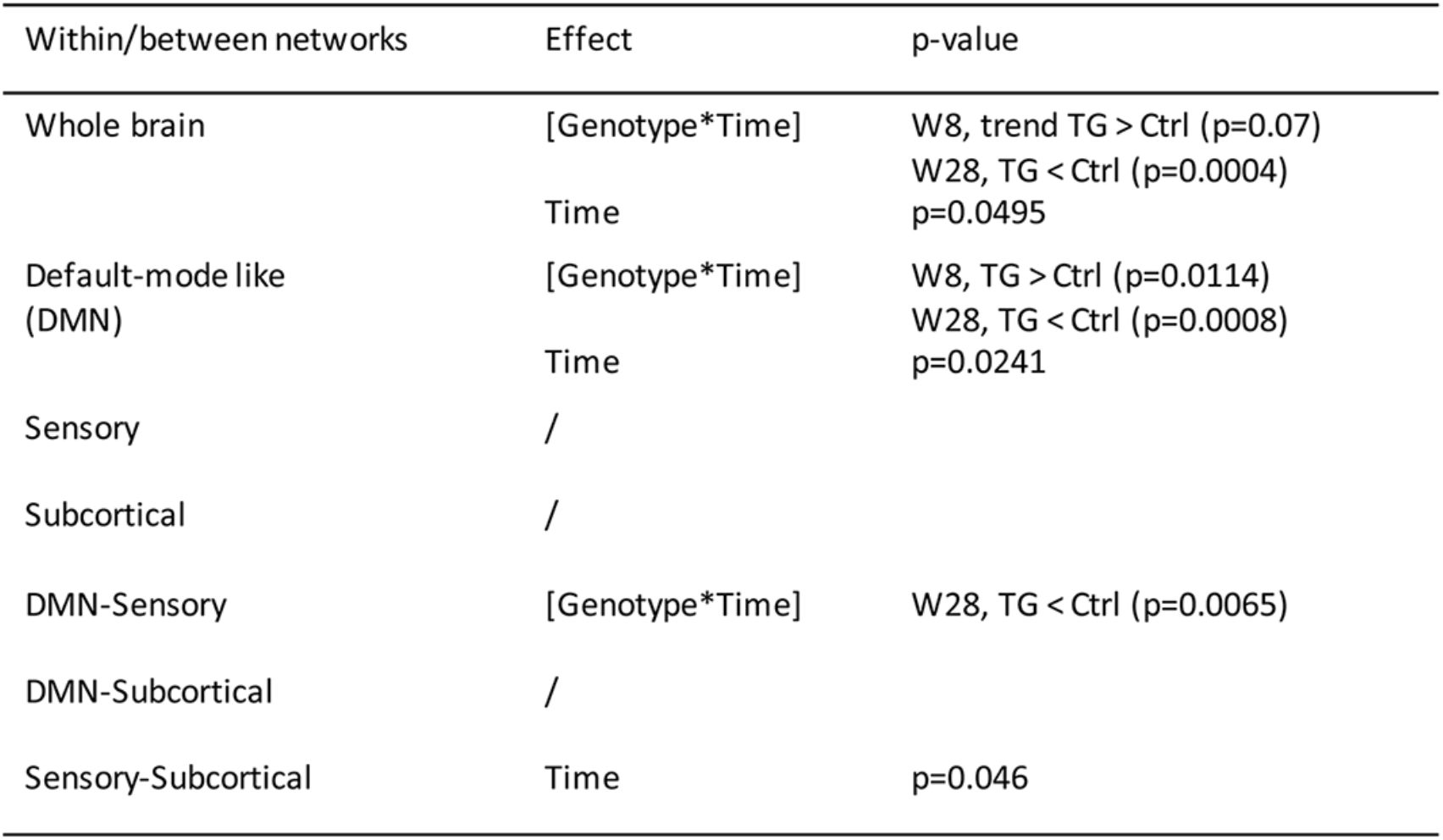
Table summarizing the outcome of the linear mixed model statistical analyses of the different networks. Abbreviations: see **Table 1**.

Interestingly, a significant hypersynchronisation was observed in the DMN of TG animals (p=0.0114) relative to controls at 8w post-DOX treatment and FC was then decreased in a similar way to the whole brain analysis leading to a significant hyposynchronisation relative to controls at 28w post-DOX treatment (p=0.0008, Figure 2B). On the contrary, no significant differences were found within the sensory and subcortical networks at any time point (Figure 2 C-D) indicating that the effect was more specific to the DMN.

### TG mice showed altered FC within the DMN

As the biggest differences were found in the DMN, we further investigated this network, in order to assess which regions were more affected. Figure 3 illustrates the difference of the average FC (TG - Ctrl) of a specific region within the DMN over time by spherical nodes localized in that brain region and scaled according to the amplitude of the FC difference (see also Additional Figure 3 for time-courses and Table 3 for a summary of the statistical analysis).

**Figure 3.**
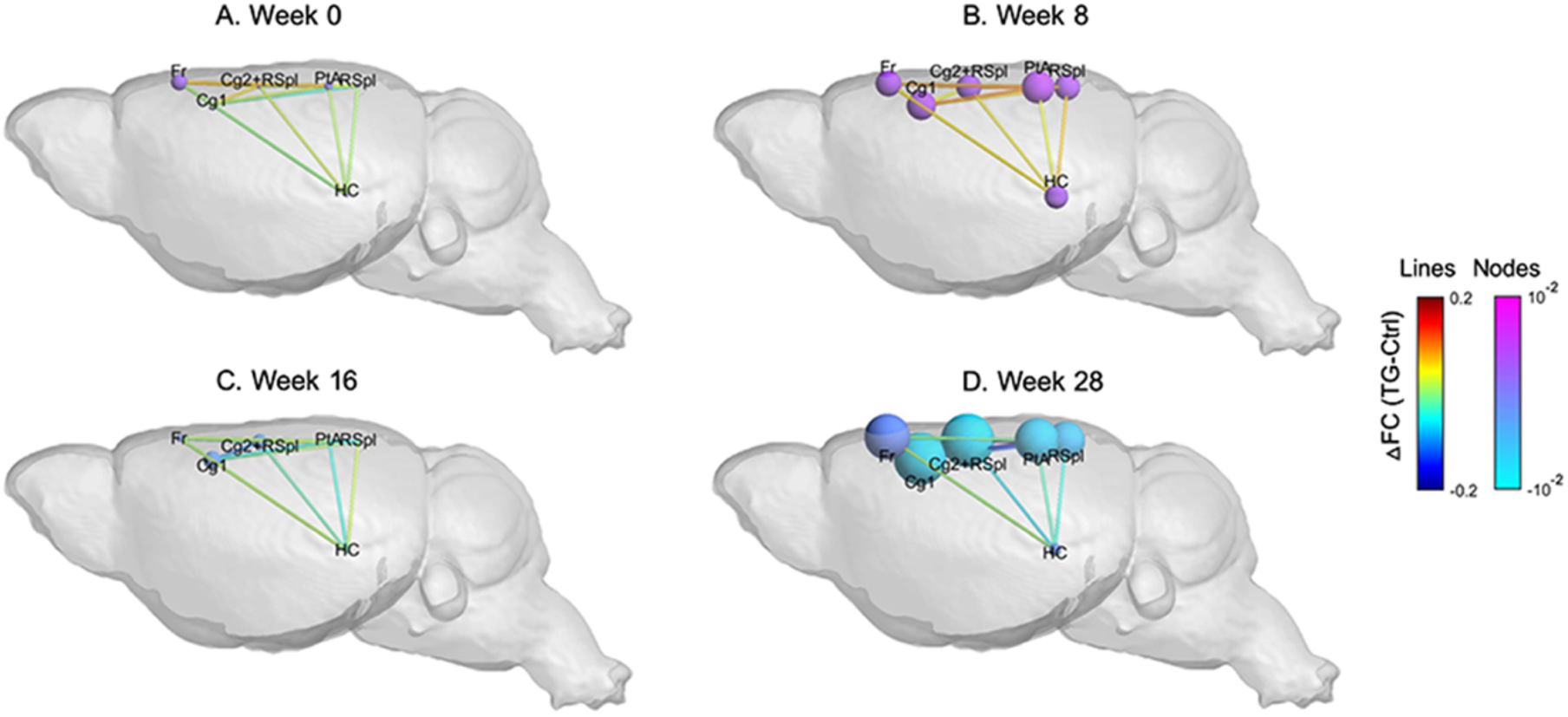
Aberrant FC in the default mode-like network in the Tet-Off APP mice. The brain schematics illustrate the difference in FC within (nodes) and between (lines) regions in the DMN over time, overlaid on a 3D anatomical template surface. The inter-node FC difference is represented by the lines, with the colour scale illustrating the actual FC difference between Tet-Off APP and Ctrl (TG-Ctrl), with orange indicating a stronger connection in the TG mice. The intra-node FC difference is described by larger or smaller spheres, with pink indicating a larger node FC within the DMN in Tet-Off APP mice compared to age-matched Ctrl mice.

**Additional Figure 3.**
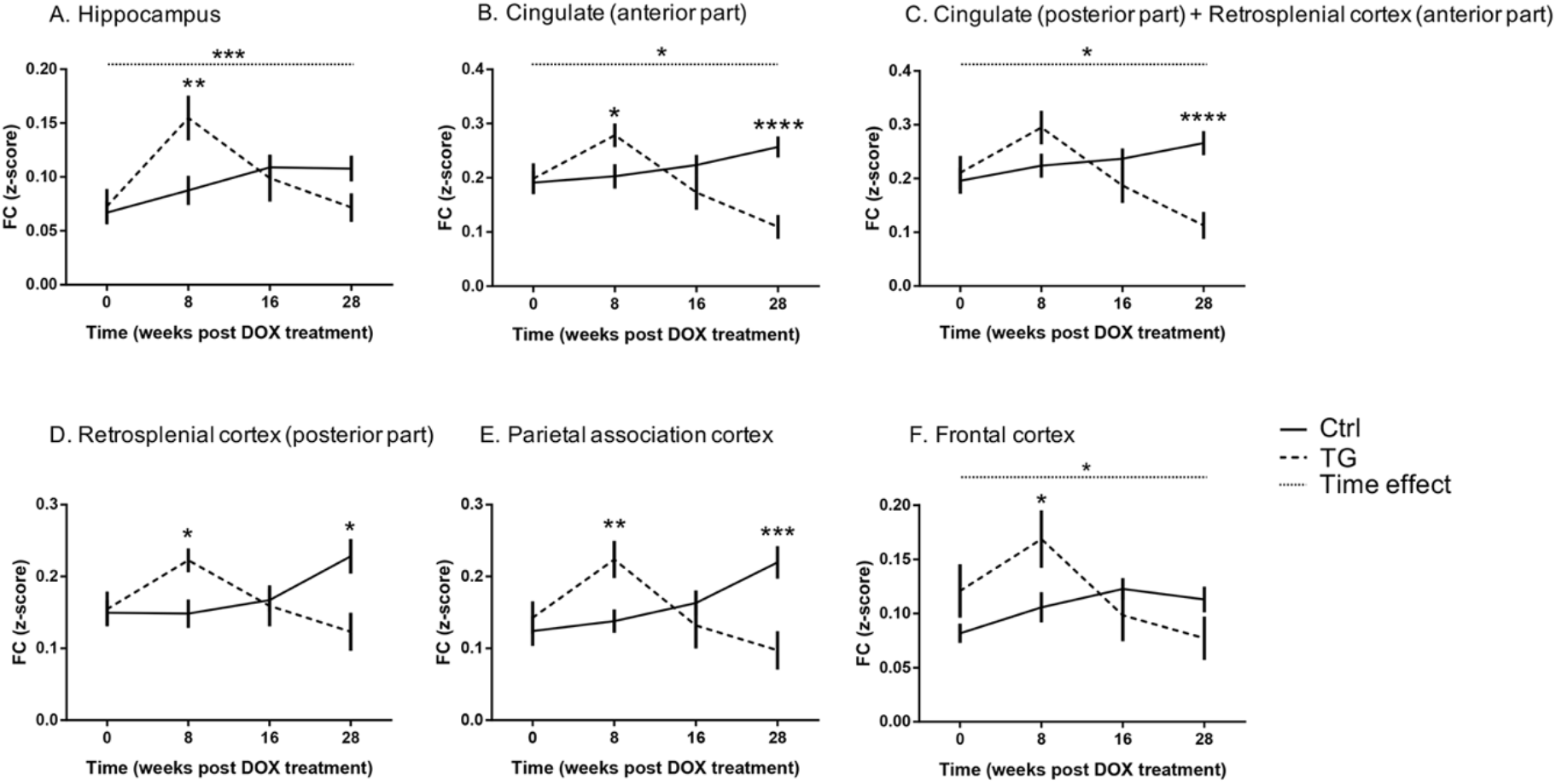
Time evolution of FC in regions of the DMN network. Each graph shows the FC over time of a specific region. Results are shown as mean ± SEM. Significant [Genotype*Time] interaction and Time effect are indicated with stars and dotted line, respectively. *p<0.05; **p<0.01; ***p<0.001; ****p<0.0001.

**Table 3.**
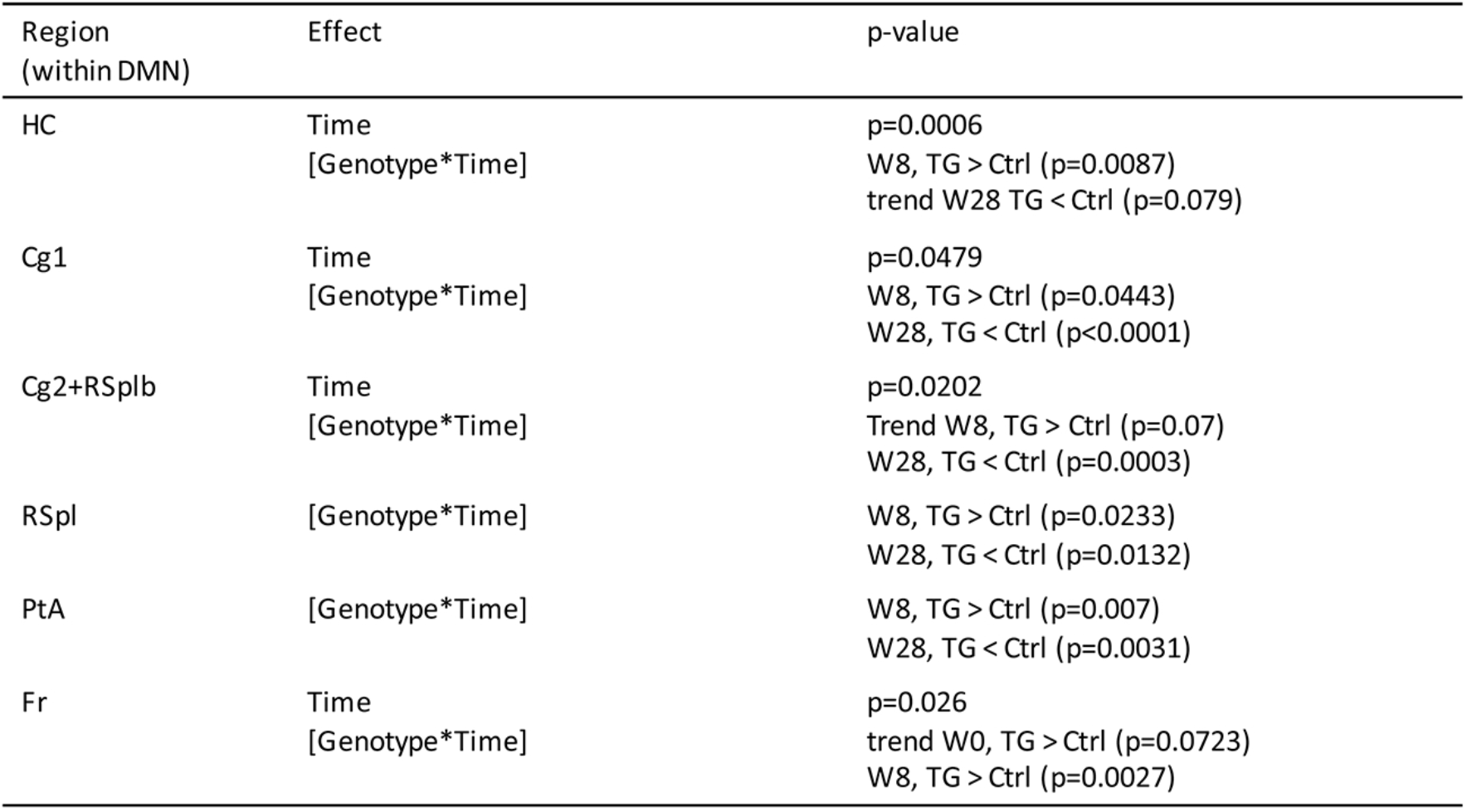
Table summarizing the outcome of the linear mixed model statistical analyses of the different regions of the DMN. Abbreviations: see **Table 1**.

At week 0 post DOX treatment, most of the regions showed small differences (Figure 3A). At 8w post DOX treatment, the FC differences were larger, reflecting an increased FC in the TG animals for all regions (Figure 3B). More specifically, the TG mice showed a significant increase in FC compared to the controls in the hippocampus (p=0.0087), the anterior part of the cingulate (p=0.0443), in the posterior part of the retrosplenial cortex (p=0.0233), in the parietal association cortex (p=0.007) and the frontal cortex (p=0.0027). At 16w post DOX treatment, all the FC differences were reduced (around 0), with the TG mice showing lower FC for all regions (Figure 3C). At the last time point, 28w post DOX treatment, the difference in FC were remarkably large, indicating an overall reduced FC for the TG group (Figure 3D). Indeed, the TG mice showed a significant decreased FC in the anterior part of the cingulate (p<0.0001), the posterior part of the cingulate cortex-anterior part of the retrosplenial cortex (p=0.0003), the posterior part of the retrosplenial cortex (p=0.0132) and the parietal association cortex (p=0.0031, Additional Figure 3).

### TG mice showed an altered inter-region FC in the DMN

After the assessment of the average FC of each region within the DMN, we evaluated how the different regions are functionally connected to each other. Therefore, we looked at the difference in FC between each pair of regions in the DMN, between the TG and controls (Figure 3; lines). A summary of the statistical analyses performed on the inter-regional FC is presented in Table 4.

**Table 4.** Table summarizing the outcome of the linear mixed model statistical analyses of the different regions of the DMN (after FDR correction for multiple comparison of p<0.05). Abbreviations: see **Table 1**.

| Connection<br>(within DMN) | Effect | p-value |
| --- | --- | --- |
| Cg1-Cg2+RSplb | [Genotype*Time] | W28, TG < Ctrl (p=0.0001) |
| Cg1-Fr | [Genotype*Time] | W8, TG > Ctrl (p=0.0473)<br>W28, TG < Ctrl (p=0.0221) |
| Cg1-PtA | [Genotype*Time] | W8, TG > Ctrl (p=0.0393)<br>W28, TG < Ctrl (p=0.0005) |
| Cg1-RSpl | [Genotype*Time] | W8, TG > Ctrl (p=0.0482)<br>W28, TG < Ctrl (p=0.0096) |
| Cg2+RSplb-Fr | [Genotype*Time] | Trend W8 TG > Ctrl (p=0.0745)<br>Trend W16 TG < Ctrl (p=0.0638) |
| Cg2+RSplb-PtA | [Genotype*Time] | W28, TG < Ctrl (p=0.0002) |
| RSpl-PtA | [Genotype*Time] | Trend W8 TG > Ctrl (p=0.0514)<br>W28 TG < Ctrl (p=0.0224) |
| HC-Cg1 | Time | p=0.0212 |
| HC-Cg2+RSplb | Time | p=0.0212 |
| HC-Fr | Genotype | p=0.0349 |
| PtA-Fr | Genotype | p=0.025 |

The inter-regional FC differences followed a similar time pattern as the average FC per region. We observed an overall increased FC in the TG animals compared to the controls at 8w post DOX treatment (Figure 3B), followed by a decreased FC between all pairs of regions 28w post DOX treatment (Figure 3D). In fact, some connections demonstrated a statistically significant main effect of time or main effect of genotype (Figure 4A).

**Figure 4.**
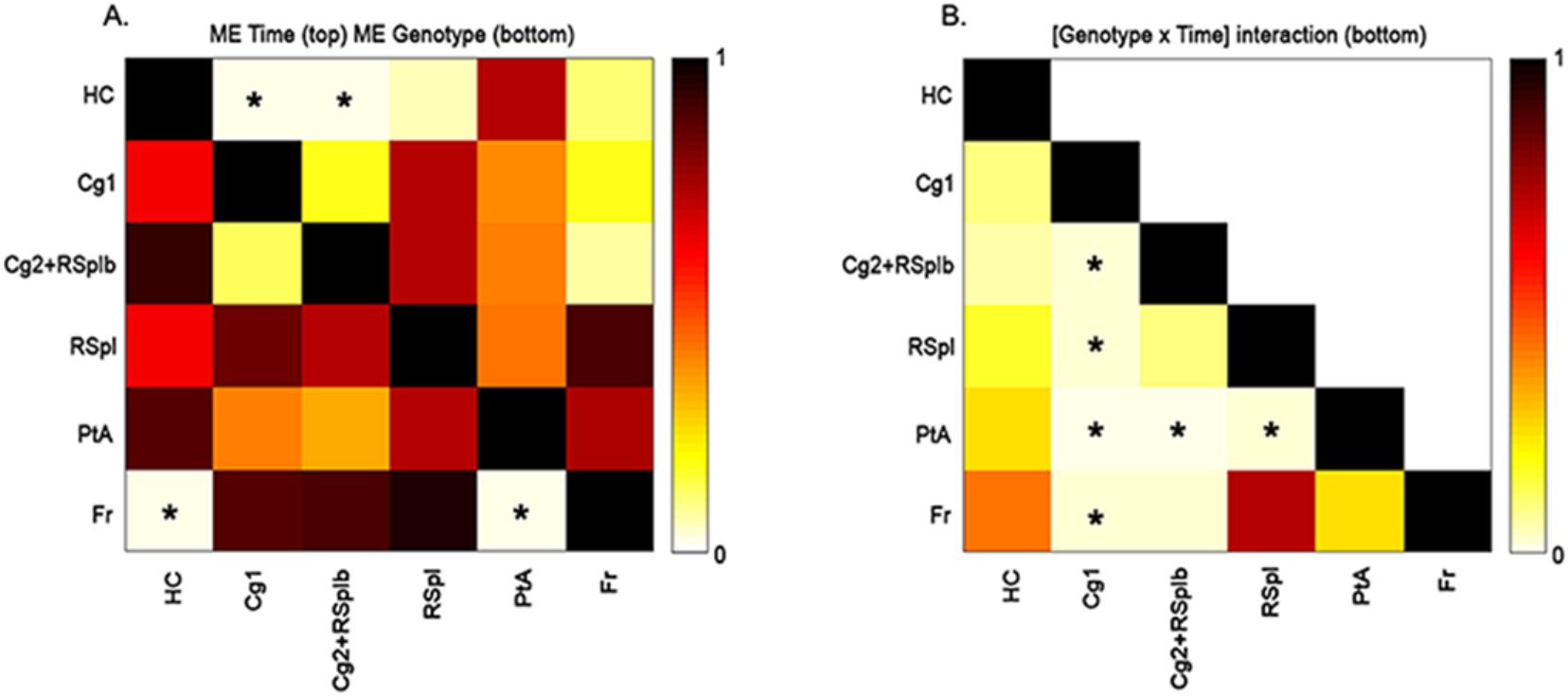
Outcome of the linear mixed model statistical analysis per connection. **A-B**. The matrices show the p-values after FDR correction for multiple comparisons of p<0.05 of the main effects (**A**) of time (sub-diagonal) and genotype (supra-diagonal) and of the [Genotype*Time] interaction (**B**). Each square indicates the p-value between each pair of ROIs. The colour scale represents the p-value power with light yellow indicating a low p-value and dark brown indicating higher p-values. The significant p-values are indicated with a star. Abbreviations: see **Table 1**.

More specifically, the connections between the hippocampus and the anterior part of the cingulate cortex and between the hippocampus and the posterior part of the cingulate-anterior part of the retrosplenial cortex were significantly different over time (p=0.0212). Moreover, we found two connections which presented a statistically significant main effect of genotype: the connection between the frontal cortex and the hippocampus (p=0.0349) and the connection between the frontal cortex and the parietal association cortex (p=0.025). Furthermore, in total six connections showed a statistically significant interaction [Genotype*Time] (Figure 4B). At 8w post DOX treatment, the TG mice showed greater FC for some connections, between the anterior part of the cingulate cortex with the frontal cortex (p=0.0473), between the anterior part of the cingulate cortex with the parietal association cortex (p=0.0393) and between the anterior part of the cingulate cortex with the posterior part of retrosplenial cortex (p=0.0482). In addition, two connections showed a trend, the posterior part of the cingulate-anterior part of the retrosplenial cortex with the frontal cortex (p=0.0745) and the posterior part of the retrosplenial cortex with the parietal association cortex (p=0.0514). On the other hand, at 28w post DOX treatment, the TG mice showed lower inter-regional FC compared to the control group, in particular for the following connections: the anterior part of the cingulate with the posterior part of the cingulate-anterior part of the retrosplenial cortex (p=0.0001), the anterior part of the cingulate with the frontal cortex (p=0.0221), the anterior part of the cingulate with the parietal cortex (p=0.0005), the anterior part of the cingulate with the posterior part of the retrosplenial cortex (p=0.0096), the posterior part of the cingulate-anterior part of the retrosplenial cortex with the parietal association cortex (p=0.0002) and the posterior part of the retrosplenial cortex with the parietal association cortex (p=0.0224).

### Soluble Aβ levels significantly increased 8w post DOX treatment in the TG mice

In order to confirm the presence and the increase of sAβ in the TG mice, an ELISA test was performed at 8w post DOX, revealing significant higher levels of total sAβ in the TG mice (p=0.0238). Indeed, the sAβ levels were about 20-fold higher compared to Ctrl ([sAβ]_TG_=(700±3) pg/ml vs ([sAβ]_Ctrl_=(36±6) pg/ml). Moreover, a representative Thioflavin-S staining performed at this time point confirmed the absence of Aβ plaque load in the TG mouse (Figure 5). However, at 28w post DOX treatment, the TG mouse showed numerous Aβ plaques in the cortical and hippocampal regions (Figure 5) compared to an age-matched Ctrl.

**Figure 5.**
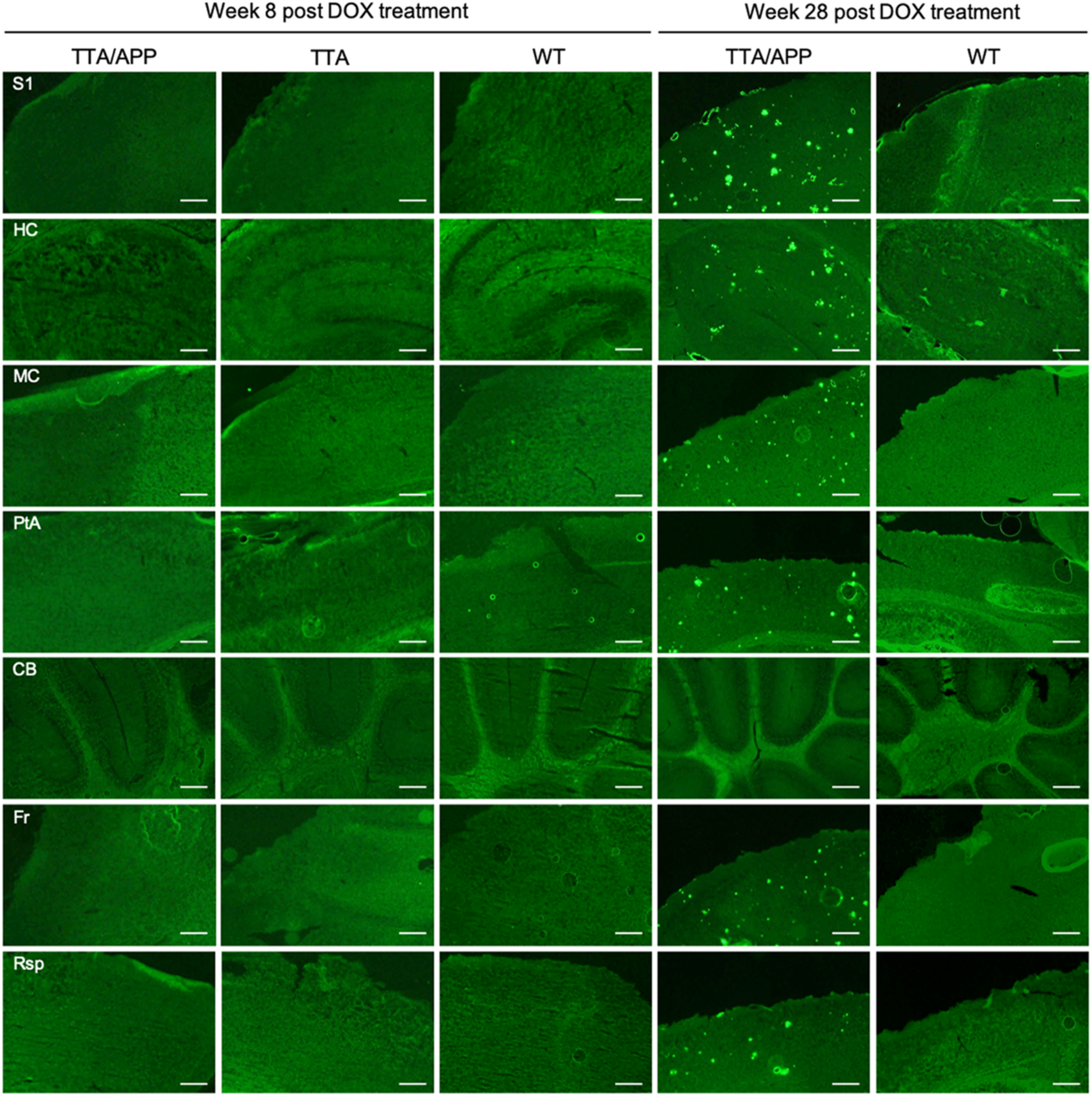
Representative images of ex vivo amyloid pathology. Overview showing the absence of amyloid β plaque load (green) in the TG mice at 8w post DOX treatment and their presence in the cortical and hippocampal regions 28w post DOX treatment, as compared to age-matched control mice. Scale bar, 275 μm.

### Hypersynchronisation of brain networks was not associated with astrogliosis or microgliosis in the TG mice

In order to check whether the aberrant FC in the TG group was associated with inflammatory responses (generally observed in AD pathology), microgliosis and astrogliosis were evaluated by staining for Iba1+ and GFAP+, respectively (Figure 6). We did not observe obvious difference in accumulation of astrocyte or activation of microglia in the TG mice compared to age-matched control animals at 8w post DOX. However, at 28w post DOX, reactive astrocytes and microglia were clearly present surrounding the plaques.

**Figure 6.**
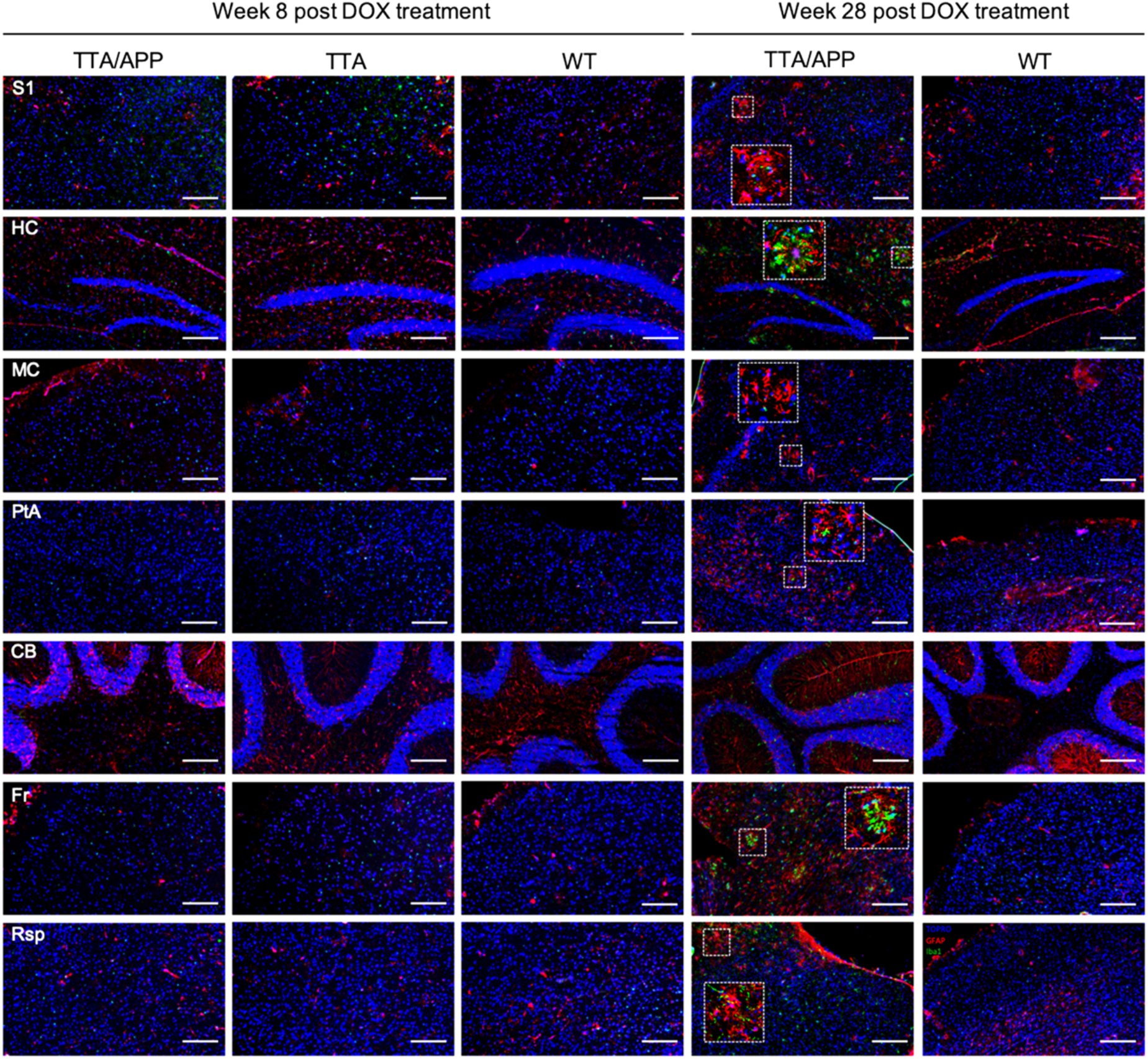
Evaluation of inflammatory responses in the Tet-Off APP mice. At 8w post DOX treatment, no obvious any sign of microgliosis (Iba1, red) or astrogliosis (GFAP, green) was found in the TG mice and control littermates. At 28w post DOX treatment, the TG mouse showed an accumulation of astrocytes and microglia surrounding the plaques, as compared to an age-matched control mouse (nuclear counterstain TOPRO in blue). Scale bar, 200 μm.

## DISCUSSION

This study aimed to address the effects of sAβ on the neuronal networks in mature-onset amyloidosis TG mice. By inducing APP overexpression only once the main developmental processes have occurred, we avoided the possible impact of protein overexpression on the brain during postnatal development and its potential consequences later on, once the brain is mature. We conjecture, that this model can thus provide valuable information on the role of amyloid pathology in sAD, for which this cascade is usually triggered well post brain development. Even though the TG mice are bearing mutations of fAD, the clear advantage of this model over other developmental-onset AD mouse models is the possibility for controlling transgene APP expression in time. Despite this clear advantage, one should keep in mind that administration of DOX could possibly alter and inhibit microglia activation, which can contribute in the mice’s phenotype. Against this hypothesis, however, the TG mice developed the amyloid pathology, with microglia responses similar to other models [32]. While only resembling amyloid aspects observed in human AD pathology [3, 4], the inducible Tet-Off APP mouse model used here is highly relevant for studying Aβ-induced synaptic dysfunction on the neuronal networks in mature-onset TG mice.

Here, by controlling chimeric APP expression in the Tet-Off APP model, we could for the first time demonstrate spatiotemporal dynamics of brain networks in mature-onset mice by means of non-invasive and translational rsfMRI. Particularly, we could assess the impact of increased sAβ levels on the brain networks from the initial stages up to abundant Aβ plaque deposition, and correlate the neuronal network alterations to sAβ-induced synaptic toxicity.

Of note, at week 0 post DOX treatment, the TG mice did not show any significant difference in FC compared to the controls. While the TG animals carry genetic mutations, those manipulations had no detectable or apparent effect on the normal brain networks function during the suppression phase of APP transgene expression. However, 8w later, the TG animals showed a significant hypersynchronisation in the DMN, in which both intra- and inter(sub)network FC were significantly higher compared to the controls (Figure 2B, 3B, 4 and Additional Figure 3). This hypersynchronisation of the neuronal networks in the TG mice coincided with a significant increase of total sAβ levels in absence of plaques (Figure 5), suggesting a sAβ-induced toxicity affecting the brain networks and the communication between them before plaque deposition. Moreover, this neuronal network hypersynchronisation was not associated with any detectable inflammatory responses (Figure 6), while neuroinflammation is an important feature of human AD pathology. Thus, our current results showed an early hypersynchronisation of brain networks associated with sAβ increases similarly to previously reported studies using developmental-onset mouse models [29]. Noteworthy, the brain networks affected were strongly model-dependent, making the comparison between studies difficult. For instance, 3-month-old TG2576 mice showed early hypersynchronisation in the hippocampus, followed by a hypersynchronisation in the DMN, the thalamus, the cingulate and piriform cortices two months later. These alterations of brain networks were demonstrated to be associated with an increased ratio of excitatory/inhibitory synapses [29]. However, in PDAPP mice, an early hypersynchronisation was only found in the frontal cortex at 3m of age. The variability observed across the mouse models of amyloidosis could originate from model specific characteristics. Accordingly, in APP-overexpressing transgenic mouse models of amyloidosis, the mutant APP is either the human transgene or a chimeric mo/huAPP transgene bearing humanized Aβ domain with single or multiple fAD-associated mutations [3]. Independent of the type of mutations incorporated in the transgene, the levels and start of APP transgene overexpression might also interfere with the observed phenotype [4]. Although the role of APP is still not fully understood, APP was shown to be involved in synaptic functions and plasticity [38, 39] as well as in GABAergic neurotransmission, at pre- and postsynaptic level under normal physiological conditions [40]. On the contrary, an overexpression of APP could lead to an opposite effect, being toxic for the neurons. Furthermore, the promotor used to construct the transgene may also have an impact, inducing strong and persistent transgene expression not *per se* restricted to neurons, or to the brain, with some active *in utero*. Some of these effects have been reported in the literature, e.g. a persistent locomotor hyperactivity in adult mice [34], an overall reduction in the number of adult-generated hippocampal neurons [41], cognitive deficits and pathological features unrelated to Aβ levels [42] or an abnormal electroencephalogram (EEG) activity [43]. However, the early intra- and internetwork FC hypersynchronisation seems to be due to increased sAβ levels rather than the effect of APP overexpression, as demonstrated in the knock-in APP^NL-F/NL-F^ mouse model of amyloidosis expressing APP at physiological level, which exclude the interference of APP artefacts [44]. Interestingly, a similar phenomenon was observed in humans. Asymptomatic young adults with fAD showed greater activation in the right anterior hippocampus compared to matched controls [45]. Cognitively-intact children at genetic risk for fAD showed an early hypersynchronisation of the DMN, which coincided with significant higher plasma levels of Aβ_1-42_ [46]. Furthermore, healthy young adult APOE-ε4 carriers (known as greatest genetic risk factor for sAD) displayed increased activation in the DMN [47, 48] and in the hippocampus [49]. This hyperactivation seems to be persistent in the hippocampus of older adult APOE-ε4 carriers [50]. Interestingly, at 16w post DOX treatment, the TG mice showed similar FC values as compared to the controls. This decreased FC comparable to controls’ FC might be explained by the decreased of sAβ levels, as the Aβ plaques started to deposit around this time point as confirmed by Thioflavin-S staining of brain slices (data not shown), suggesting that the plaques might be inert, protecting the brain from sAβ toxicity, as a dynamic equilibrium might exist between toxic oligomers and inert plaques [51].

Nevertheless, the continuous production and accumulation of sAβ revealed to exacerbate its toxicity after the wide-spread deposition of amyloid plaques in cortical areas and hippocampus. Indeed, at 28w post DOX treatment, an overall hyposynchronisation within and between brain networks were observed in the TG mice, and more specifically in the DMN. These findings are consistent with previously reported studies using developmental-onset amyloidosis models in which significantly weaker intra- and internetwork FC was observed at advanced stages of Aβ pathology [27, 29, 44]. In effect, 18-month-old APP/PS1 mice (plaques deposition starting at 2m of age) showed weaker FC across the entire brain, as well as lower interhemispheric FC in the somatosensory cortex and the hippocampus at advanced stages of Aβ pathology [27]. In the knock-in APP^NL-F/NL-F^ mice (plaques deposition starting at 6m of age), a telencephalic hyposynchronized FC was found at 7m of age, as well as hyposynchronized FC between hippocampus and prefrontal cortex, striatum and hippocampus, and striatum and cingulate cortex [44]. However, the TG2576 and PDAPP mice demonstrated hyposynchronisation of brain networks before Aβ plaque deposition, which worsened with age [29]. These observations are in agreement with human data. Indeed, patients with mild cognitive impairment and AD patients demonstrated to have a consistent decreased FC in the DMN compared to healthy controls [17, 52, 53]. Interestingly, the regions prone to sAβ-induced hypersynchronisation in absence of plaques tend to deteriorate into subsequent hyposynchronisation after Aβ plaque burden. This decrease connectivity in the neuronal networks seems to be caused by a progressive sAβ-induced synaptic damage.

Since sAβ is continuously produced and amyloid plaques might also serve as a reservoir, the subsequent gradual and persistent synaptic loss reflected by first a hypersynchronisation followed by a hyposynchronisation of neuronal networks might come from a failure of the homeostatic mechanisms [54]. Indeed, the accumulation of sAβ oligomers in the extracellular space could reflect a deficiency in metabolites clearance or in vascular hemodynamic responses, as an increased cerebral blood flow seems to compensate the increased amyloid burden in cognitively normal Aβ+ elderly [55]. Thus, the synapse silencing might be a compensatory mechanism to counteract sAβ-induced hyperactivity of the neurons, leading however to memory deficits at advanced stage of AD. As a change in synaptic transmission provide a physiological substrate for cognition, it is not surprising that the DMN is vulnerable and affected by sAβ during AD progression [56]. The DMN network includes regions which are closely involved in cognitive functions, such as the hippocampus in learning and memory processing [57]. However, its capacity to neurogenesis might be limiting for the homeostatic regulatory system in stabilizing this hub of plasticity [54]. Indeed, adult hippocampal neurogenesis is sharply reduced in AD patients [58]. Thus, hippocampal networks are more prone to become disturbed early in AD than the primary sensory cortices, which remain fully functional until late stages.

While further experiments are needed to shed light on the mechanisms underlying the early hypersynchronisation of neuronal networks in AD pathology, the inducible Tet-Off APP mouse model of amyloidosis could be an interesting and useful tool when used as mature-onset. The identification of the local concentration of most toxic sAβ species would provide crucial information on the early events triggering the pathology. Overall, the evaluation of the efficacy of the clearance mechanism and vascular responses will give insight on the possible causes for sAβ accumulation.

## CONCLUSION

In the present study, our results indicate that abnormalities in neuronal network FC, associated with sAβ synaptotoxicity, also occur when APP expression is induced once the brain is mature. As the developmental influence of APP and/or sAβ has been avoided, this precludes that these deleterious effects on the neuronal networks were caused during postnatal brain development or a consequence of its high exposure during this critical period. Thus, the Tet-Off APP mouse model could be an interesting and useful tool while used as a mature-onset model to study the amyloid aspect of sAD for further understanding the AD mechanisms. Moreover, the combination of this inducible APP expression model with early non-invasive *in vivo* rsfMRI readout for sAβ synaptotoxicity sets the stage for future Aβ targeting preventative treatment studies.

## ABBREVIATIONS

(s)Aβ: (soluble) amyloid-beta
AD: Alzheimer’s disease
ANTs: Advanced Normalization Tools
APP: amyloid precursor protein
BOLD: blood oxygen level-dependent
CamKIIα: calmodulin-dependent protein kinase type II alpha chain
Ctrl: control
DEA: diethylamine
DMN: default mode (like) network
DOX: doxycycline
EEG: electroencephalography
ELISA: enzyme-linked immunosorbent assay
EPI: echo planar imaging
fAD: familial form of AD
FC: functional connectivity
GE: gradient echo
ICA: independent component analysis
NFTs: neurofibrillary tangles
PBS: Phosphate-buffered saline
PCR: polymerase chain reaction
ROI: region of interest
rsfMRI: resting state functional magnetic resonance imaging
sAD: sporadic form of AD
tetO: tetracycline-responsive
TG: Tet-Off APP
tTA: tetracycline-Transactivator
VOI: volume of interest
WT: wild type

## ETHICS DECLARATIONS

### Ethics approval and consent to participate

All procedures were performed in strict accordance with the European Directive 2010/63/EU on the protection of animals used for scientific purposes. The protocols were approved by the Committee on Animal Care and Use at the University of Antwerp, Belgium (permit number 2014–76). All animals were housed in a room in constant temperature (22±2°C) and humidity (55±10%), with a 12h light/dark cycle (light on at 07:00 AM) and free access to food and water available *ad libitum*. All efforts were made to ensure animal welfare.

### Consent for publication

Not applicable.

### Availability of data and materials

The materials used and/or analysed during the current study are available from the corresponding author on reasonable request.

## AUTHOR CONTRIBUTIONS

MV, AVDL and GAK conceived and supervise the work, designed the experiments and offered data interpretation advice. IBN performed the longitudinal imaging experiments. JD performed immunostaining and ELISA experiments with the supervision of PP. IBN carried-out data analyses, performed data interpretation and wrote the manuscript. AJK supervised aspects of the daily work and performed data interpretation. All authors read, edited and approved the final manuscript.

## ACKNOWLEDGEMENTS

The authors are indebted to Prof. Dr. Joanna L. Jankowsky (Baylor College of Medicine, Houston, Texas, United States) and Prof. Dr. JoAnne McLaurin (Sunnybrook Health Sciences Centre, Toronto, Canada) for providing the mice. We also express our gratitude to Dr. Jelle Praet and Johan Van Audekerke for their help.

